# Synaptic communication within the microcircuits of pyramidal neurons and basket cells in the mouse prefrontal cortex

**DOI:** 10.1101/2024.01.16.575904

**Authors:** Zsuzsanna Fekete, Filippo Weisz, Mária Rita Karlócai, Judit M. Veres, Tibor Andrási, Norbert Hájos

## Abstract

Basket cells are inhibitory interneurons in cortical structures with the potential to efficiently control the activity of their postsynaptic partners. Although their contribution to higher order cognitive functions associated with the medial prefrontal cortex (mPFC) relies on the characteristics of their synaptic connections, the way they are embedded into local circuits is still not fully uncovered. Here, we determined the synaptic properties of excitatory and inhibitory connections between pyramidal neurons (PNs), cholecystokinin-containing basket cells (CCKBCs) and parvalbumin-containing basket cells (PVBCs) in the mouse mPFC. By performing paired recordings, we revealed that PVBCs receive larger unitary excitatory postsynaptic currents from PNs with shorter latency and faster kinetic properties compared to events evoked in CCKBCs. Also, unitary inhibitory postsynaptic currents in PNs were more reliably evoked by PVBCs than by CCKBCs yet the former connections showed profound short-term depression. Moreover, we demonstrated that CCKBCs and PVBCs in the mPFC are mutually interconnected with each other. As alterations in PVBC function have been linked to neurological and psychiatric conditions like Alzheimer’s disease and schizophrenia and CCKBC vulnerability might play a role in mood disorders, a deeper understanding of the general features of basket cell synapses could serve as a reference point for future investigations with therapeutic objectives.

## Introduction

In the past decades, substantial effort has been made to associate different inhibitory cell types in cortical areas with specific functions in animal behavior. Research on parvalbumin-containing basket cells (PVBCs), a GABAergic cell type frequently studied in cortical structures is a great example of these efforts. The role of PVBCs has been demonstrated in generating oscillations (Sohal et al. 2009; Gulyas et al. 2010; Schlingloff et al. 2014; Stark et al. 2014), working memory (Kim et al. 2016; Lagler et al. 2016) and sensory processing (Cardin et al. 2009; Atallah et al. 2012; Bienvenu et al. 2012). Knowledge regarding the function of the other basket cell type, the cholecystokinin-expressing basket cell (CCKBC) on the other hand remains limited, although, due to their preference for innervating the perisomatic region of postsynaptic partners (Freund and Katona 2007; Armstrong and Soltesz 2012; Veres et al. 2017), both BC types are able to efficiently control the spiking of their targets (Miles et al. 1996; Cobb et al. 1995; Stark et al. 2014; Woodruff and Sah 2007, Veres et al. 2017). This might endow the two BC types with comparable significance within the balanced communication of pyramidal neurons (PNs) and inhibitory cells.

Despite the similarity of the targeted subcellular regions, the two BC types differ in several single-cell level features. PVBCs are equipped with voltage-gated Na^+^ channels showing fast inactivation alongside voltage-gated K^+^ channels, including Kv3.1 and Kv3.2 channels ensuring high-frequency firing (Rudy and McBain 2001; Hu et al. 2018). Additionally, P/Q-type voltage-gated Ca^2+^ channels enable precisely timed transmitter release from PVBC terminals (Rudy and McBain 2001; Hefft and Jonas 2005; Glickfeld and Scanziani 2006; Hu et al. 2014). CCKBCs on the other hand, according to data from other cortical regions like the hippocampus and amygdala, display regular spiking phenotype (Glickfeld and Scanziani 2006; Daw et al. 2009; Szabo et al. 2010; Barsy et al. 2017), are capable of asynchronous release (Hefft and Jonas 2005; Daw et al. 2009; Szabo et al. 2010) and express type I cannabinoid receptor (CB1) on their terminals (Katona et al. 1999; Hajos et al. 2000) through which their output is suppressed by signaling molecules released retrogradely from the postsynaptic pyramidal neurons (PNs)(Wilson and Nicoll 2001; Wilson et al. 2001). Data regarding PVBC properties in the mPFC are in line with these findings (Miyamae et al. 2017), CCKBC features, however, are yet to be determined.

The synaptic connectivity of the two BC types was also described as vastly different. In the hippocampus and amygdala, PVBCs were found to evoke inhibitory postsynaptic currents (IPSCs) in local PNs with shorter latency and faster kinetics than CCKBCs (Hefft and Jonas 2005; Barsy et al. 2017), while the amplitudes of these currents were described as comparable to (Glickfeld and Scanziani 2006; Daw et al. 2009; Veres et al. 2017) or larger (Szabo et al. 2010) than those evoked by CCKBC firing. CCKBCs on the other hand receive smaller excitatory currents from local PNs than PVBCs do (Glickfeld and Scanziani 2006; Andrasi et al. 2017). These connections, although, representing a major excitatory input to BCs (Ahrlund-Richter et al. 2019; Hafner et al. 2019), have not yet been characterized in association cortices like the mPFC.

Data linking altered PV interneuron function with neuropsychiatric disorders like schizophrenia (Hashimoto et al. 2003; Glausier et al. 2014; Gonzalez-Burgos et al. 2015) or Alzheimer’s disease (Hijazi et al. 2023; Shu et al. 2023) demonstrate how important the contribution of BCs is to healthy network operations. These findings also highlight the therapeutic potential of specifically targeting BC operation, a goal that may be fostered by thorough understanding of the synaptic interactions of BCs within the mPFC microcircuits. Therefore, in this study we set out to reveal the characteristics of synaptic transmission between PNs and BCs in the mouse prelimbic cortex (PrL) and to address the question whether the PrL is built up with synaptically interconnected populations of the two classes of BCs. By using *in vitro* whole-cell patch-clamp recordings we revealed the single-cell properties of identified BCs and the characteristics of excitatory and inhibitory synaptic transmission between PNs and BCs, as well as functional synaptic connections between the two BC types supported by anatomical data.

## Materials and Methods

### Animals

All procedures involving animals were performed according to methods approved by the Hungarian legislation (1998. XXVIII. section 243/1998, renewed in 40/2013) and institutional guidelines. All procedures were in compliance with the European convention for the protection of vertebrate animals used for experimental and other scientific purposes (Directive: 2010/63/EU). Every effort was taken to minimize animal suffering and the number of animals used. For this study, both male and female adult (P50-150) mice were used in the electrophysiological experiments from the following transgenic mouse strains: BAC-CCK-DsRed (n=28) (Máté et al., 2013), BAC-PV-eGFP (n=33) (Meyer et al. 2002) and Pvalb-IRES-Cre crossed with BAC-CCK-DsRed (n=3). For anatomical quantification, C57Bl6 (n=3) and VGAT-IRES-Cre::BAC-CCK-GFP-coIN (n=4) (Vereczki et al. 2021) adult (P49-150) mice were used.

### Experimental Design

#### Slice preparations

The brain was quickly removed from the skull following deep anesthesia induced by isoflurane and was placed into an ice-cold solution containing (in mM) 252 sucrose, 2.5 KCl, 26 NaHCO_3_, 0.5 CaCl_2_, 5 MgCl_2_, 1.25 NaH_2_PO_4_ and 10 glucose, bubbled with 95% O_2_/5% CO_2_ (carbogen gas). 200 µm-thick coronal slices were prepared with a vibratome (VT1200S, Leica Microsystems) and were incubated in an interface-type holding chamber for at least 1 hour in artificial cerebrospinal fluid (ACSF) that contained (in mM): 126 NaCl, 2.5 KCl, 1.25 NaH_2_PO_4_, 2 MgCl_2_, 2 CaCl_2_, 26 NaHCO_3_, 10 glucose, bubbled with carbogen gas, and was let to gradually cool down from 36 °C to room temperature.

#### Electrophysiology

For the recordings, slices were placed into a submerged-type of chamber and were perfused with ACSF kept at 32 °C with a flow rate of 1.5-2 ml/min. Patch-clamp recordings were performed under visual guidance of a differential interference contrast microscope (Nikon FN-1 model or BX61W Olympus upright microscope) using a 40x water dipping objective. Neurons were visualized with an sCMOS camera (Andor Technology, Belfast, UK) and fluorescent protein expression was tested with the aid of a mercury arc lamp. Patch pipettes (5-7 MΩ) were pulled with a PC-10 puller (Narishige) from borosilicate capillaries with an inner filament (thin-walled, OD 1.5). Pipettes used for patching interneurons for paired recordings with PNs were filled with an intracellular solution containing (in mM): 110 K-gluconate, 4 NaCl, 2 Mg-ATP, 20 HEPES, 0.1 EGTA, 0.3 GTP (sodium salt) and 10 phosphocreatine, adjusted to pH 7.3 using KOH with an osmolarity of 290 mOsm/L, while 10 mM GABA and 0.2% biocytin were added on the day of the experiments. The intracellular solution used for recording PNs contained (in mM): 54 K-gluconate, 4 NaCl, 56 KCl, 2 Mg-ATP, 20 HEPES, 0.1 EGTA, 0.3 GTP (sodium salt) and 10 phosphocreatine adjusted to pH 7.3 using KOH, with an osmolarity of 290 mOsm/L. This latter solution was used with an additional 0.2% biocytin and 10 mM GABA when paired recordings were performed between two interneurons. For optogenetic and pharmacological experiments the following intracellular solution was used (in mM): 60 Cs-gluconate, 80 CsCl, 1 MgCl_2_, 2 Mg-ATP, 10 HEPES, 3 NaCl, 5 QX-314-Cl adjusted to pH 7.4 using HCl with an osmolarity of 280 mOsm/L and 0.2% biocytin was added on the day of the experiments.

Whole-cell patch-clamp recordings were performed with a Multiclamp 700B amplifier (Molecular Devices, San Jose, CA, USA), low-pass filtered at 3 kHz and digitized at 10 kHz for recording firing patterns and at 50 kHz during the recording of postsynaptic currents in voltage-clamp mode. Data were recorded with Clampex 10.4 (Molecular Devices) or an in-house acquisition and stimulus software (Stimulog, courtesy of Prof. Zoltán Nusser, Institute of Experimental Medicine, Budapest, Hungary), and were analyzed with Clampfit 10.4 (Molecular Devices), EVAN 1.3 (courtesy of Prof. Istvan Mody, Department of Neurology and Physiology, University of California, Los Angeles, CA) and OriginPro 2018 (OriginLab Corp, Northampton, MA, USA).

#### Single-cell properties

The protocol used for recording firing patterns in current-clamp mode consisted of alternating 800-ms-long depolarizing and hyperpolarizing current steps with an amplitude increasing to +100 and −100 pA in 10 pA increments, then to +300 pA in 50 pA increments, and finally to 600 pA in 100 pA increments. A holding potential of −65 mV was applied during the recordings.

#### Paired recordings

Synaptic connections between interneurons and PNs were tested with 5 action potentials elicited at 33 Hz with a 5-second-long inter-stimulus interval and were recorded for analysis with an inter-stimulus interval of 20 seconds. Series resistance (Rs) of the postsynaptic cell was continuously monitored and recordings in which the Rs exceeded 20 MΩ or changed more than 20%, or in which the analyzed features of the postsynaptic response showed changes were not included in the analysis of postsynaptic currents. Due to the possibility of low initial release probability of synapses between CCKBCs (Andrási et al., 2017), 10 action potentials were elicited at 40 Hz during paired recordings to detect synaptic connections between interneurons. To eliminate any tonic activity of CB1 receptors, which may preclude identifying homotypic CCKBC connections, these pairs were recorded in the presence of CB1 receptor antagonist AM251 (1 µM). Postsynaptic cells were clamped at −65 mV. The presence of gap junctions between interneurons was tested by injecting 800-ms-long current steps into one of the recorded cells that induced hyperpolarization of at least 10 mV of amplitude while simultaneous voltage deflections were monitored in the other cell in current-clamp mode (I=0).

#### Miniature excitatory postsynaptic currents (mEPSCs)

Neurons were held at a holding potential of −65 mV in voltage-clamp mode to record mEPSCs in the presence of tetrodotoxin (TTX, 1 µM) and gabazine (5 µM). Events were analyzed by EVAN 1.3.

#### Pharmacological experiments

The presence of inhibitory inputs on PVBCs originating from CCKBCs was tested by evoking postsynaptic currents in PVBCs by using theta electrode stimulation in brain slices prepared from BAC-PV-eGFP mice. The electrode was placed into the slice approximately 200-250 µm away from the recorded PVBC, minimizing the chance of direct stimulation. The intensity of the 1-ms-long stimuli was set to evoke responses with the amplitude of at least 300 pA while neurons were held at −65 mV in voltage-clamp mode. QX-314 was intracellularly applied to eliminate action potential generation due to the outward flow of Cl^-^ ions upon the opening of GABA_A_ receptors. Blockade of excitatory postsynaptic currents was achieved by adding 2 mM kynurenic acid in the recording solution.

#### Optogenetic experiments

For testing PVBC inputs on CCKBCs, we performed whole-cell recordings simultaneously from CCKBCs and PNs in brain slices prepared from mice obtained by crossing the Pvalb-IRES-Cre with the BAC-CCK-DsRed mice. Offsprings were injected with AAV5-EF1a-DIO-hChR2-eYFP (200 nl, 3.2×10^12^ vg/ml, University of North Carolina Vector Core, catalog# 35509-AAV5) in the mPFC at P120 to enable optogenetic control of local PVBC activity. ChR2 expression was allowed for 4-6 weeks before animals were sacrificed for *in vitro* experiments. Optogenetic stimulation was achieved by 3 pulses of 5-ms-long blue LED light (447 nm, Thorlabs) at 20 Hz at 2.5 mW/mm^2^ intensity applied through a 40x objective. Recording postsynaptic currents at the same time from PNs and CCKBCs held at −65 mV proved the optical stimulation of PV+ cells to be successful in all slices.

### *Post-hoc* identification of cell types

After the recordings, slices were placed into a fixative solution containing 4% paraformaldehyde in 0.1 M phosphate buffer (pH=7.4) to enable *post-hoc* visualization of biocytin-filled interneurons by applying fluorophore-conjugated streptavidin (Cy3-SA or Alexa488-SA; 1:10,000, Sigma-Aldrich and Molecular Probes, respectively). Recorded PVBCs were distinguished from fast-spiking chandelier cells based on the morphology of their axons (Nagy-Pal et al. 2023), while CCKBCs showed strong DsRed expression and accommodating firing pattern. CB1 content of CCKBCs was tested with immunolabeling using rabbit anti-CB1 (Cayman, # 10006590) primary antibody at 1:1000 concentration, revealed with Alexa405-coupled donkey anti-rabbit secondary antibody (1:500, Jackson). Slices were mounted in Vectashiled (Vector Laboratories) and confocal images were taken using a Nikon C2 microscope using CFI Super Plan Fluor 20X objective (N.A. 0.45; z step size: 1 μm, xy: 0.31 μm/pixel) and CFI Plan Apo VC60X Oil objective for higher magnification (N.A. 1.40; z step size: 0.25 μm, xy: 0.08 μm/pixel). PNs were identified based on their slower membrane kinetics and their distinctive firing pattern characterized by regular spiking and characteristic after-hyperpolarization.

### Immunohistochemistry

For labeling putative contacts on PVBCs from CCKBCs, C57Bl6 wild type mice (n=3) were deeply anaesthetized and transcardially perfused with 4% paraformaldehyde in 0.1 M phosphate buffer (pH7.4). 80 µm coronal slices containing the mPFC were prepared with a vibratome (VT1000S, Leica Microsystems). After blocking in 10% normal donkey serum, the following mixture of primary antibodies was used (3 days: first night at room temperature, then at 4 °C): goat anti-CB1 (Frontier Institute, #CB1-Go-Af450, 1:1000), mouse anti-Gephyrin (SYSY, #147021, 1:1000) and guinea pig anti-PV (SYSY, #195004, 1:5000) in 0.1 M phosphate buffer with 2% normal donkey serum and 2% Triton-X solution. Then the following mixture of secondary antibodies was used: donkey-anti goat coupled with Alexa405, donkey-anti mouse coupled with Alexa488 and donkey-anti guinea pig coupled with Alexa647 (all Jackson, 1:500, 4 hours at room temperature). For labeling putative contacts on CCKBCs from PVBCs, mPFC slices were prepared from VGAT-IRES-Cre::BAC-CCK-GFP-coIN mice (n=4) as described above. After blocking in 10% normal donkey serum, the following mixture of primary antibodies was used (2 days: first night at room temperature, then at 4 °C): goat anti-CB1 (Frontier Institute, #CB1-Go-Af450, 1:1000), chicken anti-GFP (SYSY, #132006, 1:1000), mouse anti-Gephyrin (SYSY, #147021, 1:1000) and guinea pig anti-PV (SYSY, #195004, 1:5000) in 0.1 M phosphate buffer with 2% normal donkey serum and 2% Triton-X solution. Then the following mixture of secondary antibodies was used: donkey-anti goat coupled with Alexa405, donkey-anti chicken coupled with Alexa488, donkey-anti mouse coupled with Cy3, donkey-anti guinea pig coupled with Alexa647 (all Jackson, 1:500, 4 hours at room temperature). Slices were mounted in Vectashield, then confocal images were taken using a Nikon microscope with CFI Plan Apo VC60X Oil objective (N.A. 1.40; z step size: 0.125 μm, xy: 0.08 μm/pixel). Using 3D confocal images, interneurons within ∼30 µm depth from the surface of the slices were selected for analysis irrespective of the number of synaptic contacts on them. The surface of the somata and the putative contacts on it (i.e., where a bouton closely opposed the cell surface and a gephyrin puncta was found between them) were manually labeled and quantified with Neurolucida 10.53 software (MBF Bioscience).

### Statistical Analysis

#### Analysis and statistical tests

Reported features of the postsynaptic currents are based on the analysis of the first evoked response in each train of stimuli, elicited at least with 20 seconds of interval. After individually measuring the features of at least 15 postsynaptic events in the case of each recorded pair, statistical tests were applied on averaged data. The potency of events was calculated by excluding, while amplitude was calculated by including transmission failures. To assess statistical significance in the case of datasets with non-normal distribution based on the Shapiro-Wilk test, Mann-Whitney U-test was applied. Cumulative frequency distributions of mEPSC parameters were compared with Kolmogorov-Smirnov test. The level of significance was set to 0.05. Statistics were performed using Origin 2018.

## Results

### Contrasting active and passive membrane properties of basket cells in the PrL

One of the key factors that determine the function of a given cell type is how its inputs are integrated and converted into output activity. Therefore, we first studied the active and passive membrane properties of CCKBCs and PVBCs. Acute slices containing the PrL were prepared from BAC-CCK-DsRed or BAC-PV-eGFP mice, where CCKBCs or PVBCs express fluorescent proteins, respectively, (Nagy-Pal et al. 2023) allowing their identification prior to recordings. Following *in vitro* electrophysiological recordings, *post hoc* identification based on morphology (Nagy-Pal et al. 2023) and firing pattern ensured that only BCs were included in the study (Figure 1A). Analysis of voltage responses upon the injection of hyperpolarizing and depolarizing current steps revealed that CCKBCs had significantly larger input resistance, longer membrane time constant and more depolarized spike threshold than PVBCs, but the capacitance of the two BC types was comparable (Figure 1B, C_1_, Table 1). The narrow spikes of PVBCs were followed by significantly larger after-hyperpolarization potentials (Figure 1C_2_, Table 1), and PVBC firing pattern showed significantly less accommodation than that of CCKBCs (Figure 1B, C, Table 1), a notable difference between the two BC types also found previously in other cortical areas (Pawelzik et al. 2002; Glickfeld and Scanziani 2006; Daw et al. 2009; Szabo et al. 2010; Barsy et al. 2017). Consequently, the same amount of injected current elicited firing from PVBCs with higher frequencies compared to CCKBCs (Figure 1C_2_), eventually unveiling the characteristic fast-spiking phenotype. These diverse membrane properties suggest that signals in the two BC types are translated into distinct activity patterns and have different integration properties in the PrL as it has been first proposed in the hippocampus (Glickfeld and Scanziani 2006).

**Figure 1.**
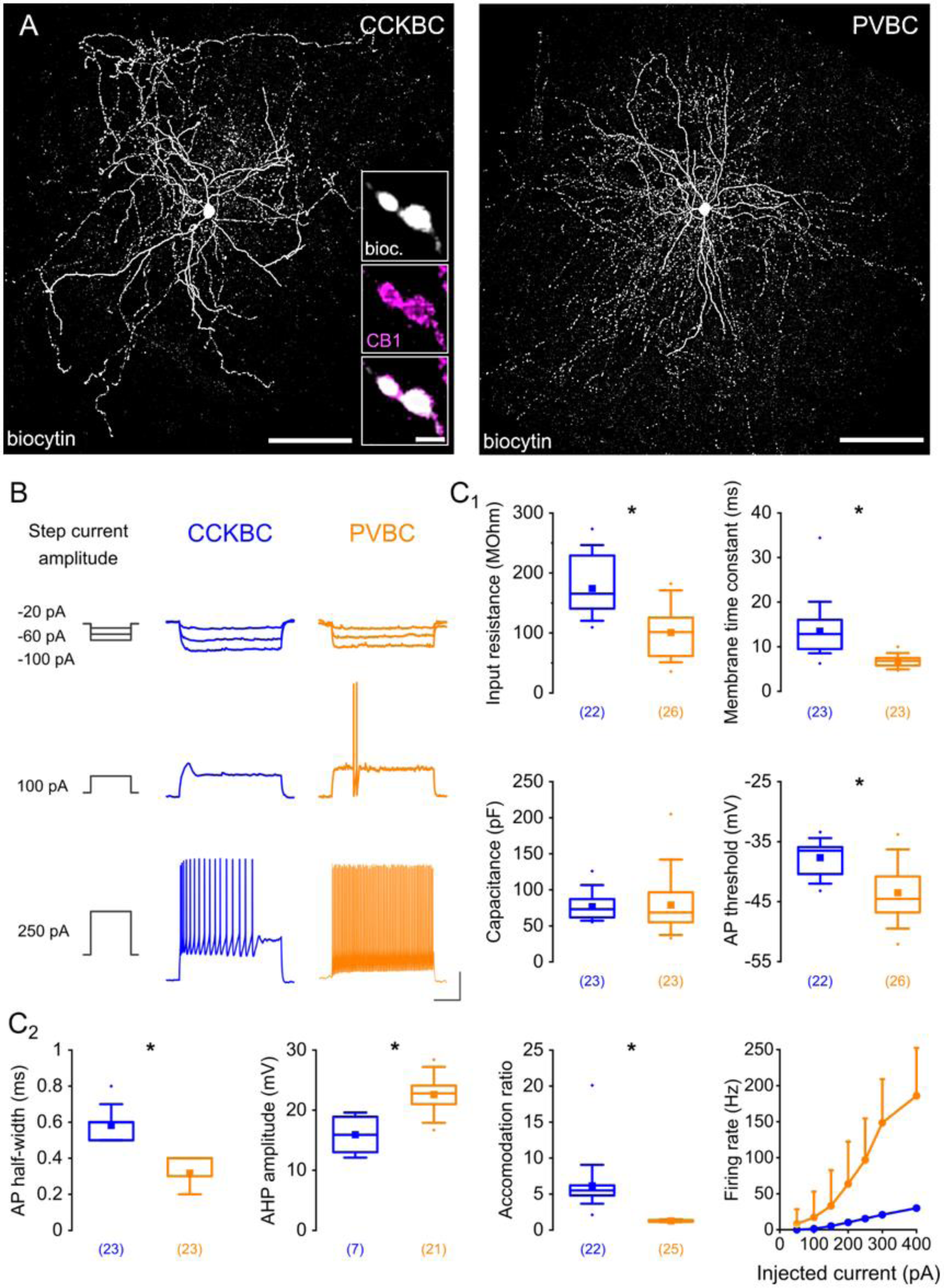
Analysis of *in vitro* recorded single-cell properties of basket cells reveals significant differences in their active and passive membrane features. (A) Confocal images taken of a biocytin-filled CCKBC and PVBC in the PrL. Inset: Axon terminals of a CCKBC co-express CB1 and biocytin. Scale bars: 100 µm and 1 µm. (B) Representative traces of basket cell responses following current step injections. Scale bar: x=200 ms, y=20 mV. (C_1-2_) Comparison of intrinsic electrophysiological properties uncovers substantial differences between the two basket cell types (M-W test, \**p* < 0.001). Boxes in this and the other figures represent the interquartile range; filled square: mean; whiskers: 5^th^ and 95^th^ percentile. Numbers in parentheses represent the number of analyzed recordings. For details see Table 1.

**Table 1.** Data are presented as the median with the 1st and 3rd quartiles in parentheses. Synaptic jitter in synaptic transmission was defined by SD of the latency values.

| Parameters |  | CCKBC |  | PVBC |  | P-value | Statistical test |
| --- | --- | --- | --- | --- | --- | --- | --- |
|  |  | median (1st and 3rd quartile) | n | median (1st and 3rd quartile) | n |  |  |
| Single-cell properties | Input resistance (MΩ) | 165.5 (140.6, 229.1) | 23 | 101.5 (61.55, 125.5) | 23 | < 0.001 | MW |
|  | MTC (ms) | 12.85 (9.49, 16.04) | 23 | 6.83 (5.78, 7.47) | 23 | < 0.001 | MW |
|  | Capacitance (pF) | 73.11 (61.63, 87.13) | 23 | 68.58 (54.97, 96.53) | 23 | 0.37 | MW |
|  | AP threshold (mV) | -36.5 (-40.4, -35.9) | 22 | -44.55 (-46.8, -40.8) | 26 | < 0.001 | MW |
|  | AP half-width (ms) | 0.6 (0.5, 0.6) | 23 | 0.3 (0.3, 0.4) | 23 | <0.001 | MW |
|  | AHP amplitude | 15.9 (13, 18.9) | 7 | 22.8 (21, 24.1) | 21 | <0.001 | MW |
|  | Accommodation ratio | 5.51 (4.81, 6.2) | 22 | 1.27 (1.15, 1.38) | 25 | < 0.001 | MW |
|  | Max. firing rate (Hz) | 41.9 (31.3, 48.8) | 22 | 208.8 (171.3, 228.8) | 23 | <0.001 | MW |
| uEPSC | Potency (pA) | 17.27 (16.06, 19.8) | 13 | 60.56 (35.04, 84.46) | 20 | < 0.001 | MW |
|  | Amplitude (pA) | 15.09 (13.02, 17.4) | 13 | 50.89 (27.95, 84.46) | 20 | < 0.001 | MW |
|  | Failure rate | 0.12 (0.03, 0.19) | 13 | 0.06 (0, 0.16) | 20 | 0.523 | MW |
|  | Latency (ms) | 1.23 (0.84, 1.8) | 12 | 0.64 (0.48, 0.82) | 18 | < 0.001 | MW |
|  | Synaptic jitter | 0.25 (0.17, 0.36) | 12 | 0.13 (0.1, 0.19) | 20 | 0.003 | MW |
|  | RT 10-90% (ms) | 0.79 (0.41, 0.81) | 13 | 0.3 (0.21, 0.35) | 20 | < 0.001 | MW |
|  | T50 (ms) | 2.86 (1.68, 3.5) | 12 | 1.29 (0.84, 1.71) | 18 | < 0.001 | MW |
| mEPSC | Instantaneous rate (s<sup>-1</sup>) | 10.09 (5, 23.16) | 5 | 20.5 (9.28, 45.91) | 5 | 0.0076 | KS |
|  | Amplitude (pA) | 15.67 (12.96, 20.06) | 5 | 27.38 (18.68, 43.41) | 5 | < 0.001 | KS |
|  | RT 20-80% (ms) | 0.44 (0.26, 0.67) | 5 | 0.21 (0.13, 0.36) | 5 | 0.048 | KS |
|  | Decay time constant (ms) | 1.99 (1.96, 2.5) | 5 | 1.33 (1.29, 1.41) | 5 | 0.022 | MW |
| uIPSC | Potency (pA) | 29.83 (16.65, 60.18) | 19 | 38.21 (29.96, 120.62) | 21 | 0.062 | MW |
|  | Amplitude (pA) | 18.71 (9.39, 38.3) | 19 | 38.21 (29.8, 120.62) | 21 | 0.004 | MW |
|  | Failure rate | 0.35 (0.08, 0.5) | 19 | 0 (0, 0.06) | 21 | < 0.001 | MW |
|  | Latency (ms) | 1.38 (0.98, 1.71) | 19 | 0.71 (0.63, 0.83) | 21 | < 0.001 | MW |
|  | Synaptic jitter | 0.14 (0.11, 0.21) | 19 | 0.08 (0.06, 0.11) | 21 | <0.001 | MW |
|  | RT 10-90% (ms) | 0.61 (0.53, 0.75) | 17 | 0.58 (0.47, 0.66) | 21 | 0.086 | MW |
|  | T50 (ms) | 4.55 (3.34, 5.55) | 16 | 4.08 (3.31, 4.95) | 21 | 0.56 | MW |
| Homotypic connections | Potency (pA) | 26.7 (21.07, 37.53) | 4 | 33.28 (28, 116.85) | 6 | 0.241 | MW |
|  | Failure rate | 0.62 (0.5, 0.65) | 5 | 0.03 (0, 0.09) | 6 | 0.013 | MW |
|  | Latency (ms) | 1.02 (0.9, 1.36) | 4 | 0.64 (0.44, 1.06) | 6 | 0.166 | MW |

AP, action potential; AHP, afterhyperpolarization; KS, Kolmogorov-Smirnov test; mEPSC, miniature excitatory postsynaptic current; MTC, membrane time constant; MW, Mann-Whitney test; RT, rise time, uEPSC, unitary excitatory postsynaptic current; uIPSC, unitary inhibitory postsynaptic current.

### Local PNs evoke larger uEPSCs in PVBCs with short latency and fast kinetic properties

To determine how the two BC types are embedded into the local excitatory networks, we investigated the excitatory connections between PNs and BCs by performing paired recordings and analyzed the properties of the unitary excitatory postsynaptic currents (uEPSCs) evoked by the 1^st^ action potential in each spike train (Figure 2A, B, Table 1). Our recordings revealed that uEPSCs evoked in PVBCs have significantly larger amplitude than those evoked in CCKBCs (Figure 2C, Table 1). Moreover, uEPSCs in PVBCs followed the presynaptic action potentials with significantly shorter latency and less synaptic jitter (spike-to-spike variability) than those in CCKBCs (Figure 2C, Table 1), suggesting that over the course of the first milliseconds after the activation of PNs, local excitatory signals reach different BC populations. Besides this temporal preference for exciting PVBCs, release from excitatory synapses to CCKBCs or PVBCs were found to be similarly successful. Kinetic properties of uEPSCs recorded from CCKBCs, however, were significantly slower both in terms of their 10-90% rise time and half-width of events, the time span measured at the half amplitude (T50) (Figure 2C, Table 1).

**Figure 2.**
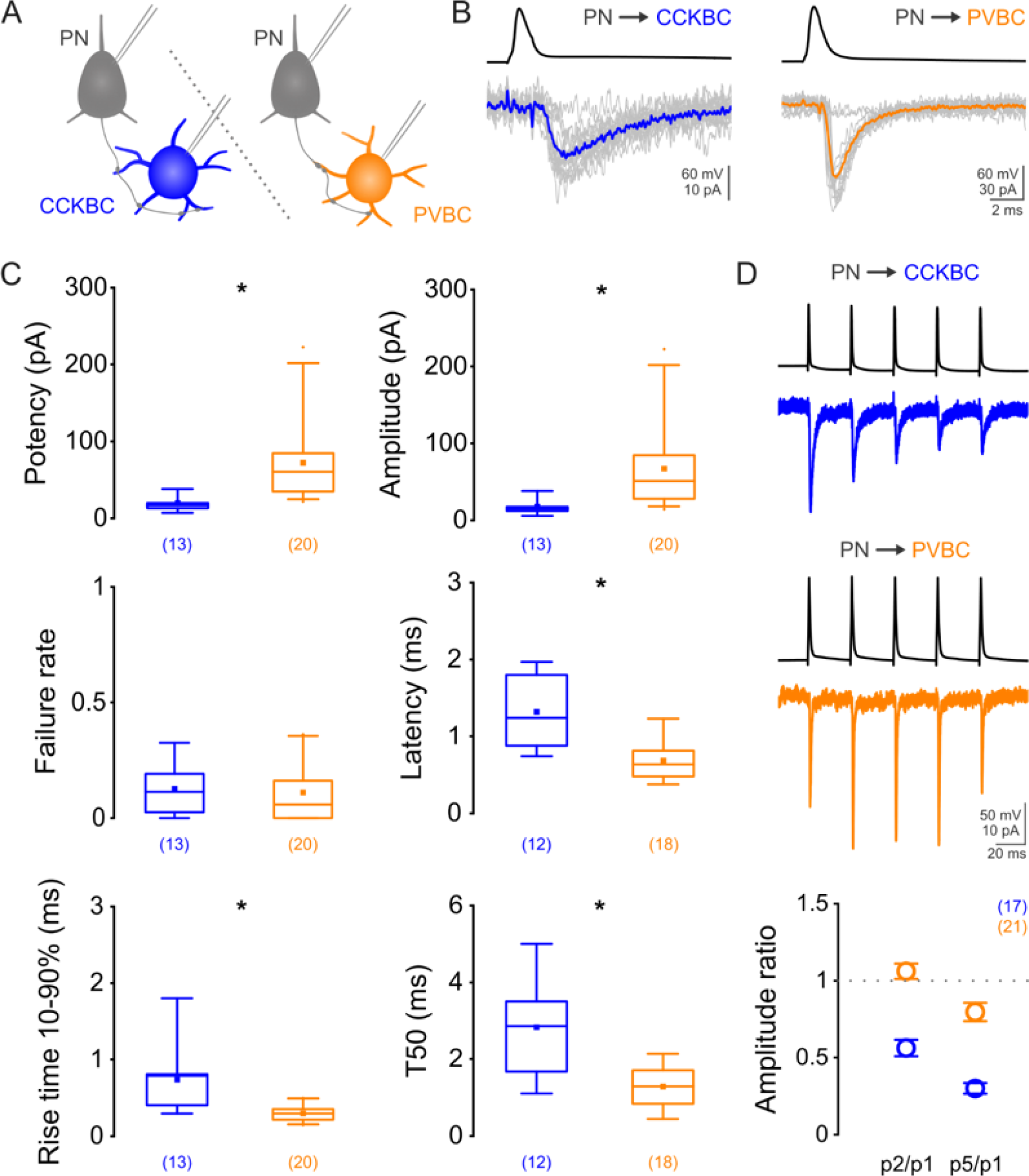
Paired recordings between PNs and BCs unveil larger unitary excitatory postsynaptic currents (uEPSCs) in PVBCs exhibiting faster kinetics than uEPSCs in CCKBCs. (A) Schematic illustration of the experiment. (B) Representative traces of the first evoked action potential (AP) in a train of 5 APs in PNs (top) and the postsynaptic uEPSCs recorded in BCs (bottom). Fifteen consecutive traces in gray, average in color. (C) Comparison of the main properties of the first uEPSCs. For details see Table 1. (D) Representative traces of synapse type-dependent short-term plasticity revealed by 5 evoked APs in the presynaptic cell (top, middle). Ratio of the amplitude of the 2nd and 1st (2/1) or the 5th and 1st (5/1) uEPSCs summarizes the short-term dynamics of the excitatory connections between PNs and BCs (bottom). Data presented as mean ± SEM. Numbers in parentheses represent the number of analyzed recordings.

Synaptic connections are characterized by diverse dynamics at the short-term time scale that are dependent on the pre- and postsynaptic partners (Ali et al. 1998; Reyes et al. 1998). To unravel whether excitatory synapses on BCs become potentiated or depressed during sustained presynaptic activation we delivered trains of square pulses to evoke 5 action potentials at 33 Hz and compared the amplitude of the postsynaptic currents recorded in BCs (Figure 2D). The amplitude of uEPSCs in PVBCs received from local PNs did not decrease considerably and remained in the 80-106% range of the first uEPSC on average (2^nd^ uEPSC: 106.08±4.96% and 5^th^ uEPSC: 79.64±5.86%). Conversely, CCKBCs were found to be innervated by depressing excitatory synapses, as the amplitude of the 2^nd^ and 5^th^ uEPSCs decreased on average to 56.23±5.31% and 30.02±3.5% of the first uEPSC, respectively. Taken together, our data suggest that the excitation generated within the prelimbic networks distinguishes between the two types of BCs in a number of aspects: PVBCs receive larger and faster synaptic excitation that is maintained if repetitive firing in PNs is induced, whereas the local PN population gives rise to smaller and slower postsynaptic events in CCKBCs that show prominent depression when trains of action potentials are evoked in PNs.

### Kinetic properties of mEPSCs

To address whether the different kinetic properties of uEPSCs from neighboring PNs were a general feature of excitation arriving onto BCs in the PrL, we recorded miniature excitatory postsynaptic currents (mEPSCs) in the two BC types in the presence of tetrodotoxin (TTX, 1 µM) and gabazine (5 µM) to block voltage-gated Na^+^ channels and GABA_A_ receptors, respectively (Figure 3A, B). Comparison of the recorded events revealed that PVBCs received mEPSCs at a higher rate and these events had larger peak amplitudes than those arriving on CCKBCs (Figure 3C, Table 1). The rise time 20-80% of miniature events on CCKBCs were slower, similarly to the unitary events evoked in paired recordings (Figure 3C, Table 1). Furthermore, peak scaled averages also demonstrate the slower decay of mEPSCs recorded in CCKBCs (Figure 3D), confirmed by the decay time constant assessed on the averaged events in case of each recorded cell (Figure 3E, Table 1). These results suggest that the differential kinetic properties of excitatory postsynaptic events obtained in paired recordings are not due to sampling biases in our data but reflect the distinct features of local excitatory inputs on the two BC types.

**Figure 3.**
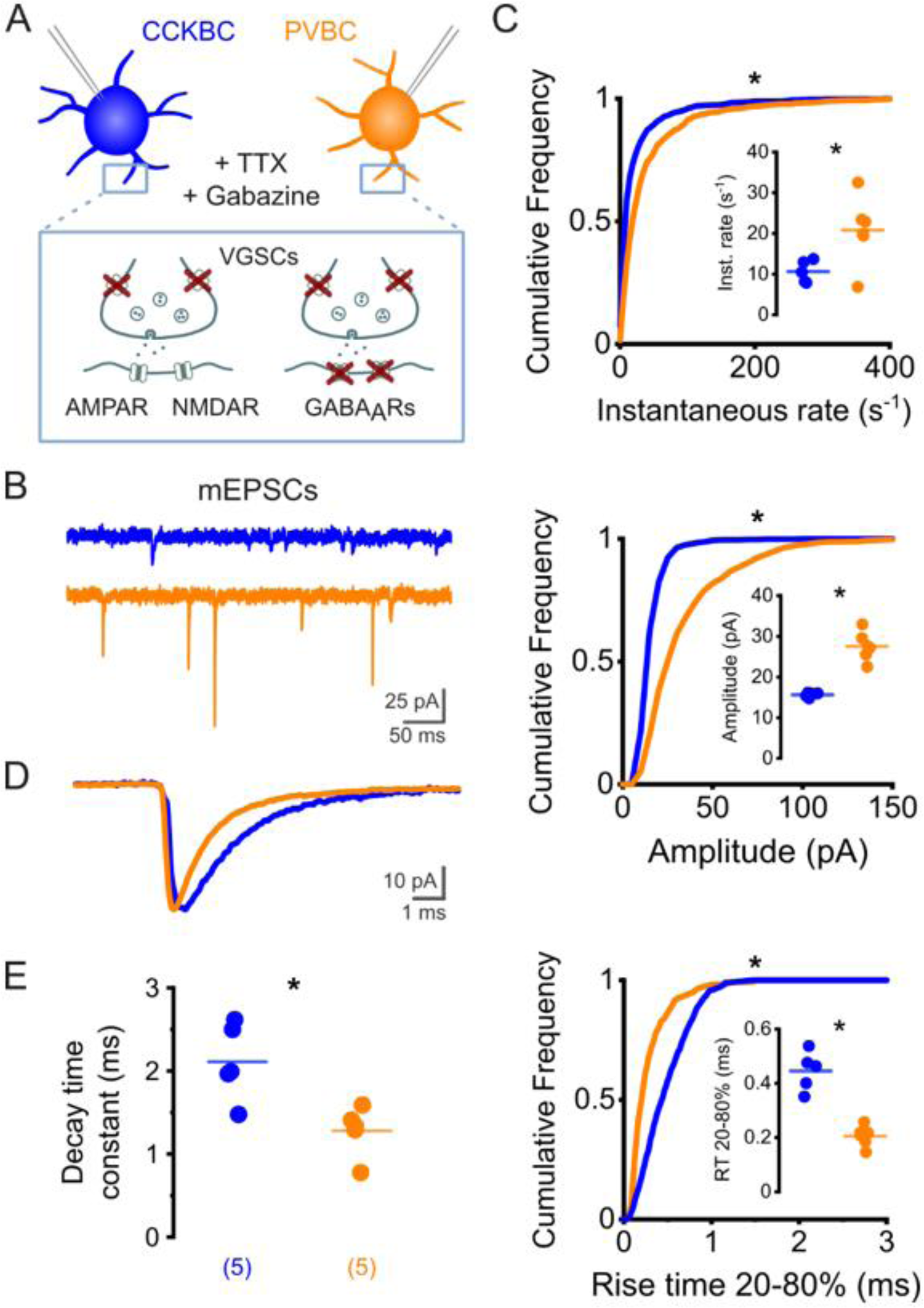
Differences in the kinetics of mEPSCs recorded in BCs resemble the uEPSCs evoked in paired recordings. (A) mEPSCs recorded in BCs in the presence of TTX (1 µM) and gabazine (5 µM) to block voltage-gated sodium channels (VGSCs) and GABA_A_ receptors (GABA_A_Rs). (B) Representative traces of recordings obtained in the two BC types. (C) Cumulative frequency plots showing that PVBCs receive miniature events with higher instantaneous rate (top), amplitude (middle) and exhibit faster rise time kinetics (bottom). Insets: median values of mEPSC parameters obtained in each cell (n = 5 for both cell types). (D) Normalized averages of mEPSCs recorded in CCKBCs and PVBCs (n = 5 for both cell type) demonstrate the slower decay kinetics of mEPSCs recorded in CCKBCs. (E) mEPSCs in CCKBCs on average are characterized by longer decay time constants. Data points represent decay time constants measured on the averages of mEPSCs recorded in individual cells. Numbers in parentheses represent the number of analyzed recordings.

### PVBCs innervate local PNs with reliable but depressing synapses

The influence of BCs on prelimbic processes is largely determined by their impact on postsynaptic partners. To uncover the effect of single BCs on neighboring PNs in the PrL we performed paired recordings between BCs and PNs (Figure 4A, B). Action potentials evoked in PVBCs elicited unitary inhibitory postsynaptic currents (uIPSCs) in PNs with significantly larger amplitude than those evoked by CCKBC spikes (Figure 4C, Table 1). While PVBC synapses were highly reliable, almost every third action potential in CCKBCs failed to evoke a uIPSC in PNs, leading to a significant difference between the failure rates of the two synapse types. Notably, postsynaptic events upon PVBC discharges showed significantly shorter latency and less synaptic jitter than those following CCKBC activation. Nonetheless, the two BC types evoke postsynaptic events in PNs with similar kinetics both in terms of their 10-90% rise time and half-width (T50) (Table 1).

**Figure 4.**
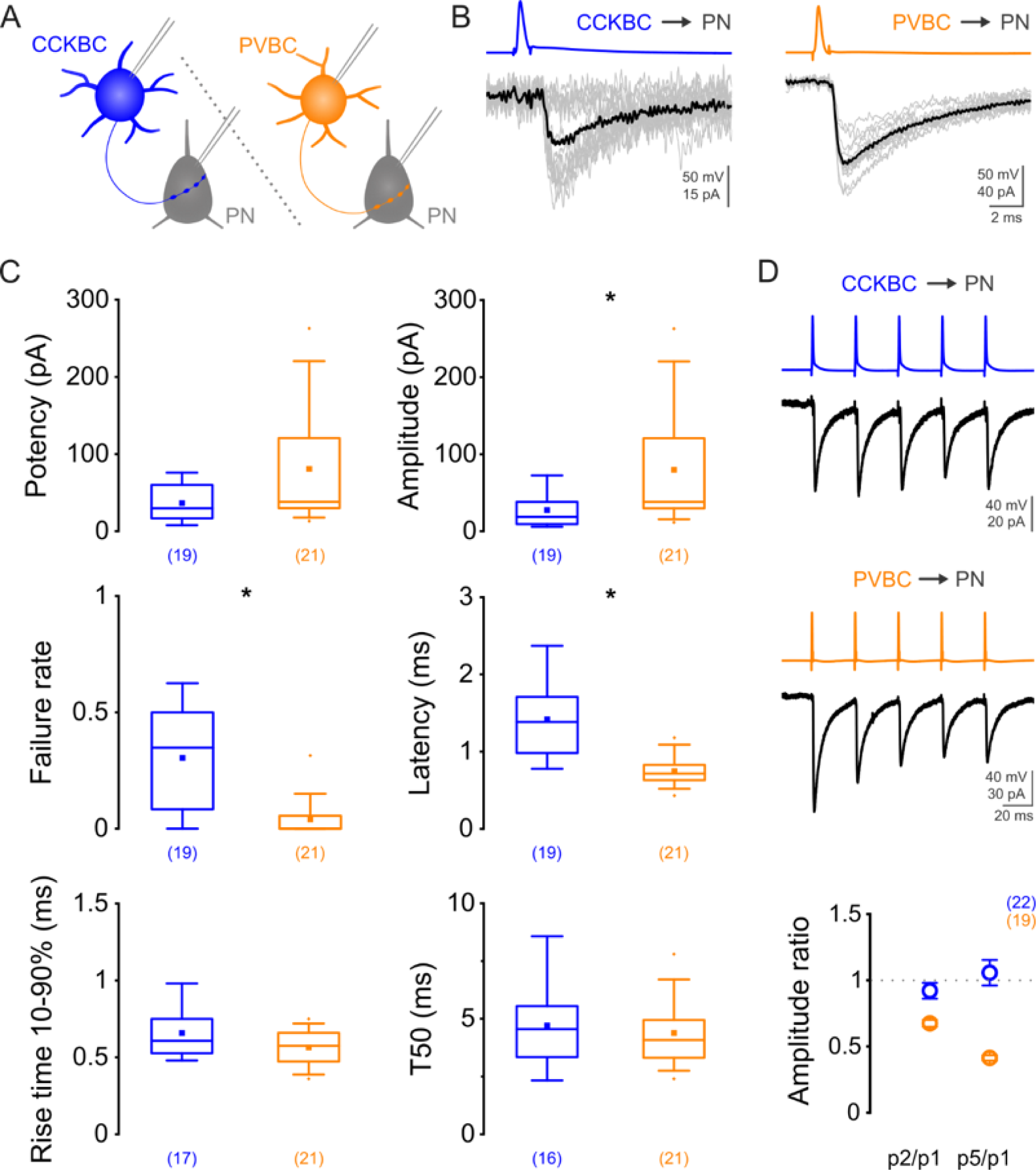
Paired recordings between BCs and PNs reveal the strong but depressing nature of synaptic inhibition arriving from PVBCs compared to uIPSCs evoked by CCKBCs. (A) Schematic illustration of the experiment. (B) Representative traces of the first presynaptic AP in a train of 5 APs evoked in BCs (top) and the postsynaptic uIPSCs recorded in PNs (bottom). Fifteen consecutive traces in gray, average in black. (C) Comparison of the main properties of the first uIPSCs. For details see Table 1. (D) Representative traces of synapse type-dependent short-term plasticity revealed by eliciting 5 APs in the presynaptic cell (top, middle). Ratio of the amplitude of the 2nd and 1st (2/1) or the 5th and 1st (5/1) uIPSC summarizes the short-term dynamics of the inhibitory connections between BCs and PNs (bottom). Data presented as mean ± SEM. Numbers in parentheses represent the number of analyzed recordings.

Investigation of the short-term dynamics in these inhibitory synapses with the same protocol described above revealed that synapses arriving from PVBCs onto PNs showed significant short-term depression, a phenomenon previously observed in other cortical regions as well (Galarreta and Hestrin 1998; Hefft and Jonas 2005; Szabo et al. 2010; Barsy et al. 2017) (Figure 4D). On the other hand, uIPSCs evoked by CCKBCs became neither potentiated nor depressed (Figure 4D) (Hefft and Jonas 2005; Galarreta et al. 2008). Taken together, our results indicate that PVBC spikes reliably evoke larger postsynaptic currents than CCKBCs in PNs, but in case of prolonged BC activation, PNs can receive a steadier level of inhibition from CCKBCs.

### Basket cells form chemical synapses and gap junctions with their own cell type

Electrical synapses between neurons of the same inhibitory cell type are generally common in cortical circuits (Galarreta and Hestrin 1999; Gibson et al. 1999; Tamas et al. 2000; Andrasi et al. 2017), while the preference of inhibitory neurons for establishing homotypic chemical synapses is cell type-dependent (Pfeffer et al. 2013; Tremblay et al. 2016). To reveal the extent to which BCs are synaptically interconnected and the electrophysiological properties of their contacts in the PrL, we performed paired recordings between the same type of BCs (Figure 5A). Testing electrical coupling was conducted by injecting hyperpolarizing current steps in one of the cells while recording membrane potential changes in the other cell (Figure 5B, top). Chemical synapses were tested by eliciting 10 action potentials (Figure 5B, bottom). CCKBCs were frequently coupled electrically (16/23 tests, 69.6%) but chemical connections seemed less numerous between them (9/31 tests, 29%), despite the elimination of any tonic activation of CB1 by AM251 application (1 µM). In contrast, every second recorded PVBC evoked uIPSCs in other PVBCs in paired recordings (13/25 tests, 52%) but only 25% of the tested cells were connected via gap junctions (2/8 tests).

**Figure 5.**
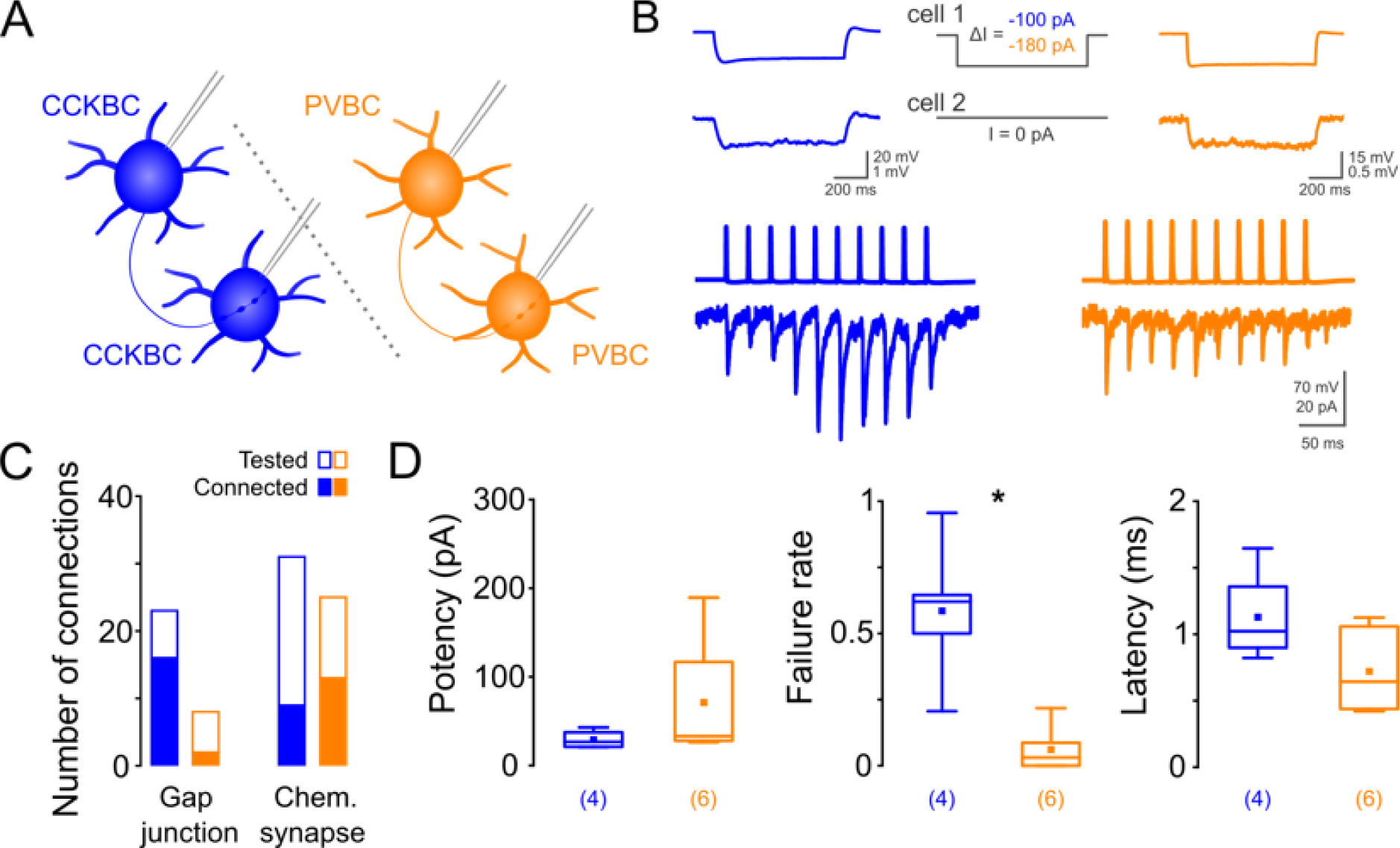
Paired recordings demonstrate that BCs innervate their own kind with chemical synapses and gap junctions as well. (A) Schematic illustration of the experiment. (B) Example traces of gap junctions revealed by injecting hyperpolarizing current steps into one of the recorded cells, averages of 50 consecutive traces (top). Example traces of recorded chemical synapses between the same type of BCs, averages of 10 consecutive traces (bottom). (C) Comparison of connection probabilities. Filled bars represent the number of connections, empty bars represent the number of tests. (D) Features of the first postsynaptic responses in homotypic connections between CCKBCs and PVBCs. For details see Table 1.

In terms of the main properties of the chemical connections, uIPSCs evoked by the first action potential tended to have larger potency in synapses between PVBCs (Figure 5D, Table 1). These homotypic PVBC connections showed short-term depression (Figure 5B), similarly to synapses between PVBCs and PNs (Figure 4D). CCKBCs on the other hand evoked larger currents in the second part of the action potential trains (Figure 5B bottom). The rate of failure for the first action potentials was significantly higher for CCKBC than for PVBC synapses (Figure 5D, Table 1) suggesting more reliable neurotransmission between the latter BCs. Although there was a tendency for uIPSCs evoked by CCKBC spikes to start off with larger latency than uIPSCs in homotypic PVBC synapses, the difference did not reach significance (p=0.17, Figure 5D, Table 1). Taken together, homotypic BC pairs display different synaptic characteristics and connectivity patterns in the PrL.

### The two basket cell types are interconnected

Previous studies have shown that the two BC types are interconnected in the hippocampus (Dudok et al. 2021) but not in the amygdala (Andrasi et al. 2017). To determine the presence or absence of functional synaptic connections on PVBCs established by CCKBCs in the PrL, we used a pharmacological approach and took advantage of CB1 expression on CCKBC axon terminals (Nagy-Pal et al. 2023, Figure 1A) that is selective for this cell type among cortical inhibitory neurons (Bodor et al. 2005). Extracellular stimulation was used to evoke IPSCs in PVBCs (Figure 6A), while glutamatergic synaptic transmission was blocked by 2 mM kynurenic acid applied in the external solution. Bath application of the CB1 agonist WIN55,212-2 (1 µM) reduced the amplitude of evoked postsynaptic currents in the recorded cells below 50% of the initial amplitude values (Figure 6B), demonstrating the presence of CB1-sensitve inputs to PVBCs. These electrophysiological data were further supported by our anatomical results showing CB1+ axonal boutons opposed to the majority of sampled PV+ somata (29 out of 33 cells, 87.9%) (Figure 6C, 6D). Immunostaining against gephyrin (the anchoring protein of GABA_A_ receptors (Sassoe-Pognetto and Fritschy 2000)) was used to ensure that only inhibitory synapses are included in the quantification. Taken together, our two independent approaches provide evidence that CB1-sensitive inputs from CCKBCs are present on PVBCs.

**Figure 6.**
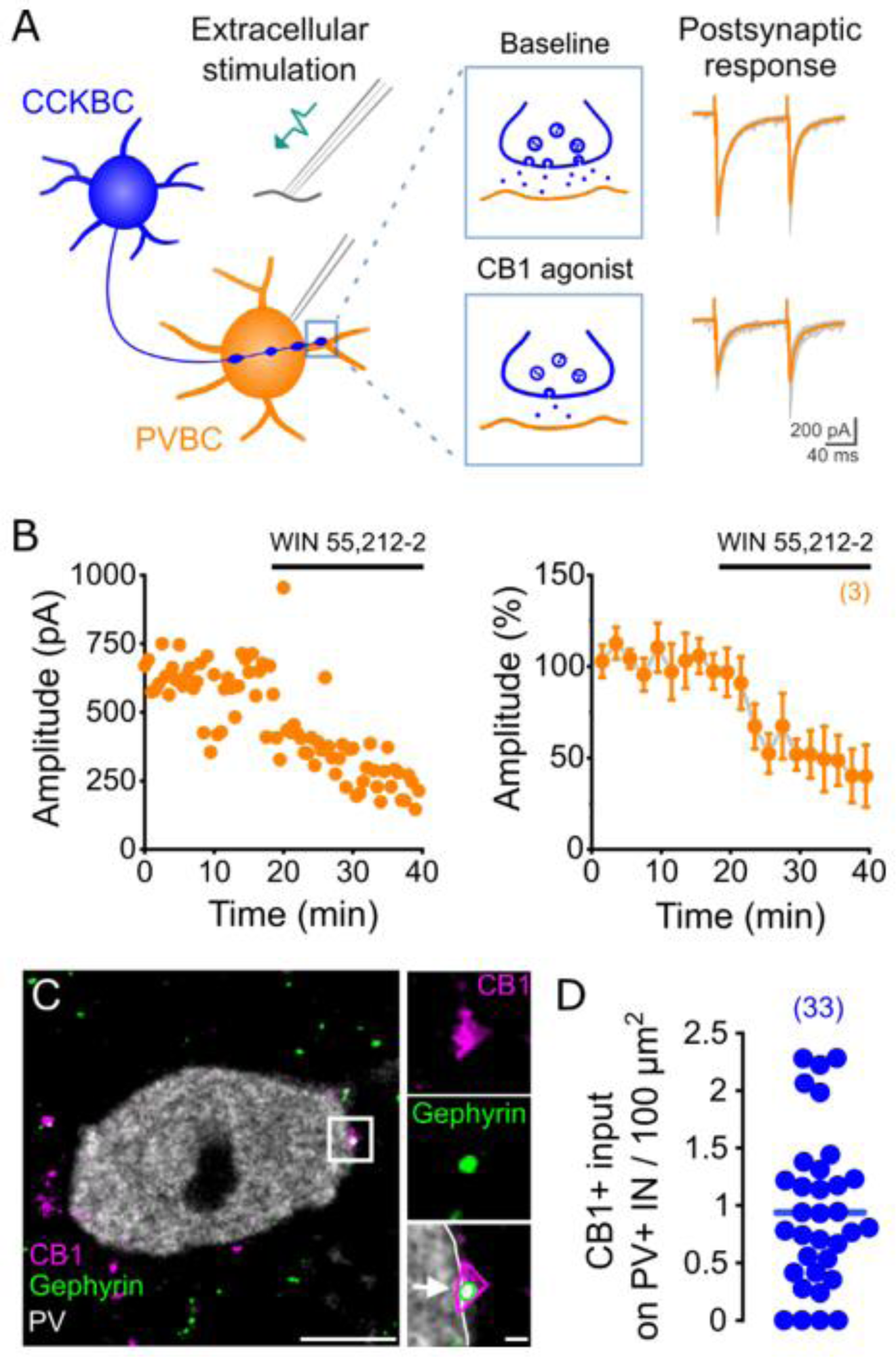
Pharmacological experiments and immunolabeling demonstrate the presence of functional CB1-positive inhibitory inputs received by PVBCs. (A) Schematic illustration of the experiment (left). Excitatory currents were blocked by 2 mM kynurenic acid in the recording solution. Representative traces of evoked IPSCs before (right, top) and after the application of CB1 agonist (right, bottom). (B) Inhibitory postsynaptic currents in a PVBC evoked by focal electrical stimulation are reduced by bath application of CB1 agonist WIN 55,212-2 (1 µM). Data from a representative experiment (left) and normalized data from n = 3 cells (right). Data points represent the mean ± SEM of 4 consecutive responses with an inter-stimulus interval of 30 seconds. (C). Confocal image of the soma of a PV+ interneuron labeled with immunostaining against PV. Inset: a terminal apposed to gephyrin labeling on the soma is immunopositive for CB1. Scale bars: 5 and 0.5 µm. (D) Quantification of the density of CB1+ terminals on PV-immunopositive somata (n=33).

To study the potential synaptic connections established by PVBCs on CCKBCs, we crossed BAC-CCK-DsRed mice with PV-Cre mice. Therefore, in offspring, we could target fluorescently labeled CCKBCs (Nagy-Pal et al. 2023) and activate PVBCs using optogenetics in the same preparations. After 4-6 weeks following the injection of AAV5-EF1-DIO-hChR2-eYFP into the PrL, we prepared acute slices and recorded from CCKBCs and PNs simultaneously in whole-cell configuration (Figure 7A). The cell types of the recorded neurons were verified by *post hoc* immunostaining against biocytin and CB1. Successful optogenetic activation of the PV+ population upon light stimulation was evident from the large IPSCs evoked in PNs (Figure 7B), while CCKBCs also received prominent inhibition without exception. The amplitude of these inhibitory currents varied between cells but the mean amplitude of IPSCs recorded from CCKBCs reached 11.1% of the IPSCs evoked simultaneously in PNs (Figure 7B). Latency of the responses measured from the beginning of light illumination was significantly larger for CCKBCs than for PNs (Figure 7B). In addition to these experiments, we also identified PV+ inhibitory synaptic contacts on the somata of CCKBCs in slices containing the PrL prepared from the offspring generated by crossing VGAT-IRES-Cre and BAC-CCK-GFP-coIN mice (Figure 7C). Immunostaining against PV and gephyrin revealed that the majority of the CCK/GFP+ somata received PV+ input (27 out of 32 cells, 84.4%) (Figure 7D). In summary, these results demonstrate the reciprocal innervation between the two BC types in the PrL.

**Figure 7.**
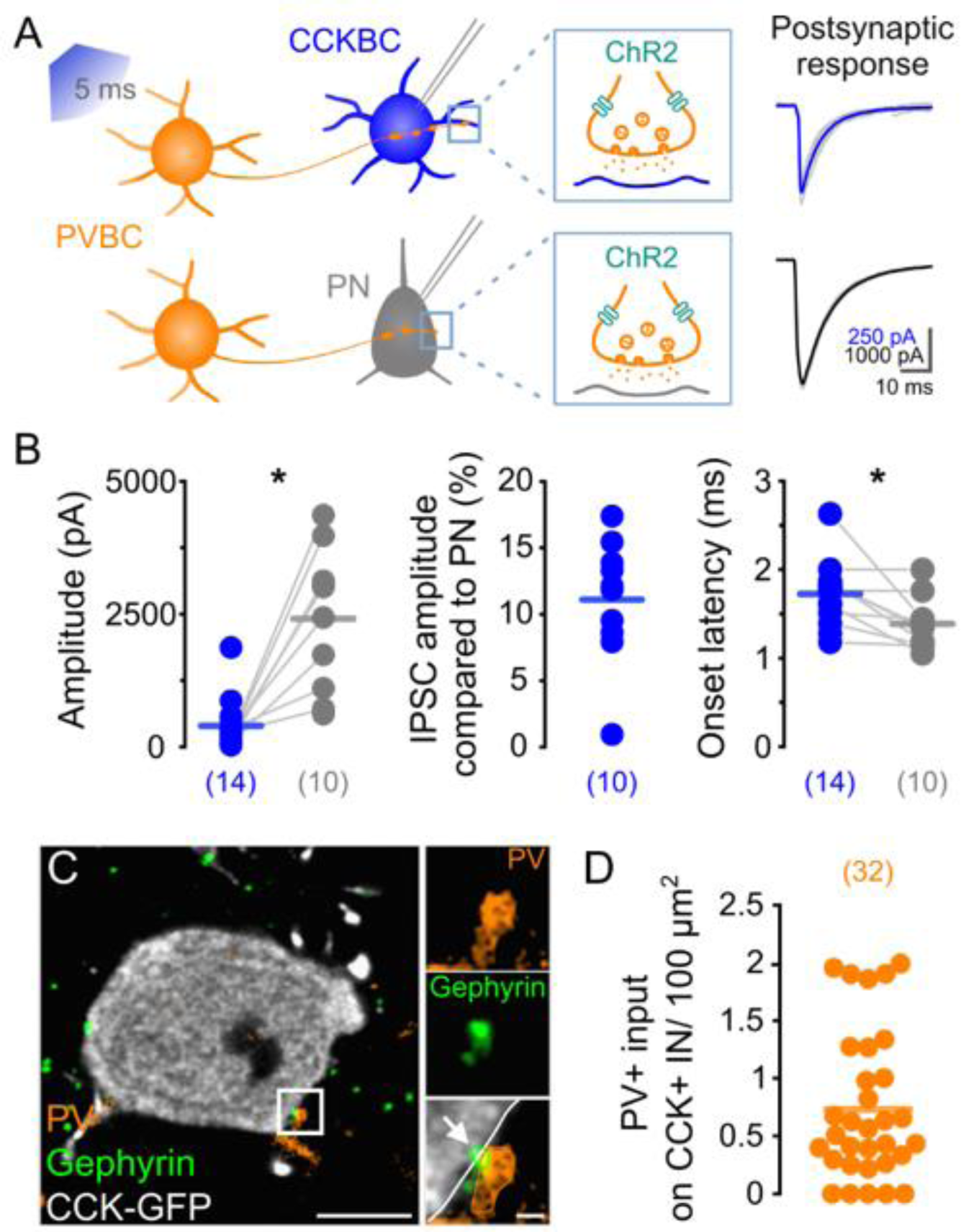
Postsynaptic responses upon light activation of PV+ cells via ChR2 together with anatomical data demonstrate that PVBCs innervate CCKBCs. (A) Schematic illustration of the experiment. Representative traces of light-evoked responses recorded simultaneously in a CCKBC (right, top) and a PN (right, bottom). Ten consecutive traces in gray, average in color. (B) Features of the light-evoked postsynaptic responses. Data points representing simultaneously obtained recordings are connected on the graph. Mean amplitude of IPSCs recorded in CCKBCs and PNs (CCKBC: 235.25 (87, 523) pA; PN: 2721.5 (1103, 3126) pA, p<0.001, MW). (left). Mean IPSC amplitude recorded in CCKBCs compared to the amplitude of IPSCs simultaneously recorded in PNs (middle). Onset latency of IPSCs (CCKBC: 1.78 (1.51, 1.84); PN: 1.35 (1.15, 1.44), p = 0.02, MW). (right). Numbers in parentheses represent the number of analyzed recordings. (C) Confocal image of the soma of a GFP-labelled CCK+ IN in the PL of a VGAT-IRES-Cre::BAC-CCK-GFPcoIN mouse. Inset: a terminal immunopositive for PV apposed to gephyrin labeling, indicating the presence of an inhibitory synapse. Scale bars: 5 and 0.5 µm. (D) Quantification of the density of PV+ terminals on CCK+ interneurons (n=32) in VGAT-IRES-Cre::BAC-CCK-GFPcoIN animals.

## Discussion

Based on our electrophysiological and anatomical data summarized in Figure 8, we determined the properties of synaptic transmission between PNs and BCs in the mouse PrL and provided evidence regarding the reciprocal connectivity among the two BC types. We showed that the membrane properties of BCs and synaptic connections with PNs display cell type specific characteristics. Our paired recordings revealed that PNs located in the PrL not only evoke larger unitary excitatory currents in PVBCs than in CCKBCs, but these currents have shorter latency. less jitter and faster kinetic parameters. Inhibition provided by PVBCs to PNs followed presynaptic activation more rapidly and reliably than the uIPSCs evoked by CCKBC firing which also tended to result in unitary currents of smaller amplitude. We provided anatomical and electrophysiological evidence that BCs innervate not only their own kind but also establish functional connections with the other BC type, a circuit motif highly relevant to our understanding of cell type function.

**Figure 8.**
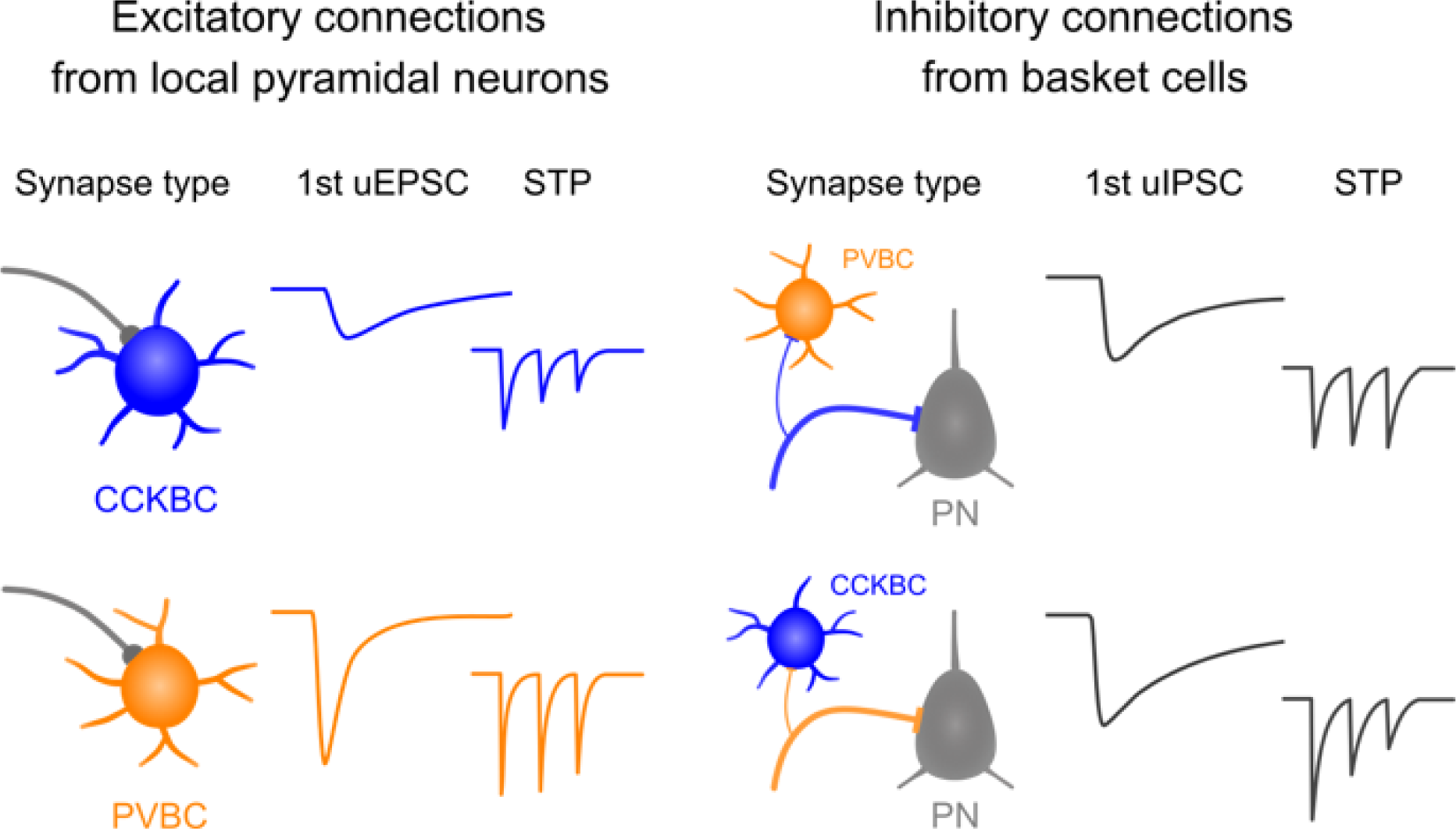
Summary of the key differences between synapses established by PNs and the two BC types within the microcircuits of the PrL. CCKBC, cholecystokinin-containing basket cell; PN, principal neuron; PVBC, parvalbumin-containing basket cell; STP, short-term plasticity; uEPSC, unitary excitatory postsynaptic current; uIPSC, unitary inhibitory postsynaptic current.

BCs have long been proposed to serve distinct circuit functions based on several differences in their single-cell features (Freund and Katona 2007). Our results regarding the contrasting input resistance, membrane time constant and accommodation ratio are in line with previous findings obtained in the hippocampus (Pawelzik et al. 2002; Glickfeld and Scanziani 2006; Cea-del Rio et al. 2010; Szabo et al. 2010; Lee et al. 2011) and basal amygdala (Andrasi et al. 2017; Barsy et al. 2017). These data provide further support to the concept that BCs are built to perform different tasks within cortical networks, despite targeting the same cellular domain in the PrL (Nagy-Pal et al. 2023).

By performing paired recordings, we showed that excitatory connections from local PNs give rise to significantly larger unitary currents in PVBCs compared to CCKBCs, in line with results obtained in the basal amygdala (Andrasi et al. 2017). Unitary EPSCs in PVBCs followed presynaptic spiking with shorter latency and less synaptic jitter than in CCKBCs, which implies a general preference for the fast and precise activation of PVBCs over the recruitment of the CCKBC populations (Glickfeld and Scanziani 2006). On the other hand, we found that transmitter release after PN activation was similarly reliable from synapses contacting either type of BCs, which is in contrast with results showing higher ratio of failure at synapses contacting CCKBCs in the basal amygdala (Andrasi et al. 2017). Given the larger input resistance of CCKBCs compared to PVBCs (Figure 1), increased transmission probability could be a way of recruiting CCKBCs more efficiently in the PrL compared to the basal amygdala. On the other hand, uEPSC on CCKBCs showed pronounced short-term depression compared to the steady uEPSCs recorded from PVBCs, a difference implying that sustained PN firing prefers to recruit PVBCs rather than CCKBCs.

Differences in the kinetics of uEPSCs were recapitulated in the rise time and decay kinetics of mEPSCs recorded in BCs. Slower rise time of miniature events in CCKBCs may suggest that excitatory synapses on this cell type are electrotonically more distant from the soma than those on PVBCs. Indeed, PVBCs receive many excitatory synaptic contacts on their somata (Gulyas et al. 1999; Hioki et al. 2013; Rovira-Esteban et al. 2020), in contrast with CCKBCs (Mátyás et al. 2004). It should be noted however, that in the basal amygdala uEPSCs in PVBCs and CCKBCs have been shown to exhibit different rise time kinetics in spite of the fact that the distance between the soma and the actual synaptic contacts were found to be similar (Andrasi et al. 2017). The contrasting decay kinetics on the other hand might be explained, at least partially, by the AMPA receptor subunit composition on PVBCs, as these cells were shown to express glutamate receptor subunit 4 (GluR4) which dictates rapid decay kinetics for EPSCs (Fuchs et al. 2007).

PVBC output synapses have long been regarded as reliable, fast transmitting sites of communication (Buhl et al. 1995; Tamas et al. 1997; Maccaferri et al. 2000; Bartos et al. 2002; Gonzalez-Burgos et al. 2005; Hefft and Jonas 2005; Glickfeld and Scanziani 2006; Galarreta et al. 2008), features that we observed in the mouse PrL as well. The first uIPSCs evoked by PVBC spikes were followed by the strong depression in amplitude, as it is typical for synapses with high release probabilities (Zucker and Regehr 2002). In contrast, CCKBCs were able to maintain a steady level of inhibition during the time window of our action potential trains, although the amplitude of these uIPSCs remained small. Our observation that decay kinetics of the BC-evoked unitary currents in PNs are comparable are in line with previous reports (Hefft and Jonas 2005; Galarreta et al. 2008; Szabo et al. 2010; Kohus et al. 2016; Andrasi et al. 2017; ^B^_arsy et al. 201_^7^; Veres et al. 2017) and the finding that hippocampal GABA_A_ receptor subunit compositions are similar in synapses on PNs facing CCKBC or PVBC axon terminals (Kerti-Szigeti and Nusser 2016). The smaller variance in decay kinetics of PVBC synapses could be, however, a necessary feature during rhythm generation, as changes in IPSC decay kinetics were shown to also impact the frequency of neuronal oscillations (Fisahn et al. 1998; Pietersen et al. 2014; Heistek et al. 2010). Apart from the similarities in uIPSC kinetics, BC output synapses are characterized by contrasting strength, precision, and reliability in the mPFC as well.

Electrical coupling and chemical synapses between inhibitory cells can serve as means of enhancing synchronous activity (Gibson et al. 1999; Tamas et al. 2000; Bennett and Zukin 2004), a circuit motif also utilized by PVBCs to aid rhythm generation in local circuits (Beierlein et al. 2000; Bartos et al. 2002). Studies show that inhibitory cells of the same class are frequently coupled (Tamas et al. 2000; Szabadics et al. 2001; Meyer et al. 2002; Galarreta et al. 2008; Andrasi et al. 2017) and our recordings obtained from CCKBCs strengthen this general concept (Iball and Ali 2011). However, PVBCs were connected to a lesser extent via gap junctions than expected based on prior studies, even though the recorded cells were located within 100 µm and innervated each other with chemical synapses in 52% of the tests. A reason behind this contradiction could be that studies reporting high connectivity rates via electrical synapses were typically obtained in juvenile animals (Coulon and Landisman 2017), while the number and strength of gap junctions between PVBCs were shown to decrease with age (Meyer et al. 2002). Our gap junction tests with PVBCs were recorded in animals older than P65 (mean age=76 days, n=4 animals), which might account for the decreased probability in electrical coupling between PVBCs. However, it cannot be ruled out that the low ratio of electrical connections between the sampled PVBCs was partly due to the considerable distance between the gap junctions and the location of the recording pipettes as these connections were shown to occur up to several hundred µm away from the somata (Fukuda et al. 2006). Chemical synapses on the other hand displayed similar tendencies in terms of their strength, reliability and latency as the synapses that contact PNs, and as in homotypic paired recordings obtained in the hippocampus (Bartos et al. 2001; Daw et al. 2009; Kohus et al. 2016).

Synaptic communication between the two BC types have only been investigated previously in a small number of studies (Karson et al. 2009; Andrasi et al. 2017; Dudok et al. 2021), despite the fact that inhibitory regulation is a major factor in the activity of interneurons as well, and consequently, in the activity of neuronal circuits (Chamberland and Topolnik 2012). According to the quantifications of immunolabeled terminals in the primary somatosensory cortex, CCKBCs indeed establish synaptic connections with PV+ cells by providing 12% of the inhibitory boutons contacting PV+ somata (Hioki et al. 2018). Moreover, paired recordings obtained in the hippocampus reported functional synapses between the two types of BCs originating from CCKBCs (Karson et al. 2009) or PVBCs (Kohus et al. 2016). On the other hand, the same method led to the conclusion that the two BC types avoid innervating each other in the basal amygdala (Andrasi et al. 2017). Our pharmacological and optogenetic approaches combined with immunocytochemical investigations provided higher throughput than dual whole-cell recordings and were able to reveal connections between BC types in the mPFC, providing a further step towards deciphering BC operation.

Our results determining the characteristics of synaptic transmission and connectivity patterns of BCs help to refine our understanding about the building blocks of circuit operation in the PrL. Deeper knowledge of connectivity in prefrontal microcircuits in healthy brain is crucial to identify potential changes when these neuronal networks operate pathologically, which typifies many psychiatric disorders, including schizophrenia, attention deficit and autism. Revealing the deviations in circuit parameters in BC networks caused by genetic and/or environmental factors may help recognizing potential targets for interventional strategies, aiming to reduce often devastating symptoms.

## Acknowledgements

We acknowledge financial support from the HUN-REN Hungarian Research Network, Hungarian Brain Research Program (2017-1.2.1-NKP-2017-00002) and National Research, Development and Innovation Office (K131893). The authors are grateful to Éva Krizsán and Erzsébet Gregori for their excellent technical assistance. We also thank P#x00E1;l VP#x00E1;gi and LP#x00E1;szló Barna, the Nikon Microscopy Center at the Institute of Experimental Medicine, Nikon Austria GmbH, and Auro-Science Consulting, Ltd., for kindly providing microscopy support.

